# Transcriptional analysis in multiple barley varieties identifies signatures of waterlogging response

**DOI:** 10.1101/2022.11.26.518028

**Authors:** Alexandra Miricescu, Ailbhe Jane Brazel, Joseph Beegan, Frank Wellmer, Emmanuelle Graciet

## Abstract

Waterlogging leads to major crop losses globally, particularly for waterlogging sensitive crops such as barley. Waterlogging reduces oxygen availability and results in additional stresses, leading to the activation of hypoxia and stress response pathways that promote plant survival. Although certain barley varieties have been shown to be more tolerant to waterlogging than others and some tolerance-related QTLs have been identified, the molecular mechanisms underlying this trait are mostly unknown. Transcriptomics approaches can provide very valuable information for our understanding of waterlogging tolerance. Here, we surveyed 21 barley varieties for the differential transcriptional activation of conserved hypoxia-response genes under waterlogging, and selected five varieties with different levels of induction of core hypoxia-response genes. We further characterized their phenotypic response to waterlogging in terms of shoot and root traits. RNA-sequencing to evaluate the genome-wide transcriptional responses to waterlogging of these selected varieties led to the identification of a set of 98 waterlogging-response genes common to the different datasets. Many of these genes are orthologs of the so-called ‘core hypoxia response genes’, thus highlighting the conservation of plant responses to waterlogging. Hierarchical clustering analysis also identified groups of genes with intrinsic differential expression between varieties prior to waterlogging stress. These genes could constitute interesting candidates to study ‘predisposition’ to waterlogging tolerance or sensitivity in barley.

## Introduction

Waterlogging (*i.e.*, the saturation of soil with water) and flooding-related stresses are the cause of major crop losses worldwide. They are predicted to worsen in some countries as a consequence of seasonal increases in rainfall resulting from global climate change (Bailey-Serres *et al*., 2012a; Bailey-Serres *et al*., 2012b; Kaur *et al*., 2020; Langan *et al*., 2022). Waterlogging and flooding alter biological, physical and chemical parameters in a plant’s environment. These combined changes affect negatively different aspects of plant growth, but at the same time, they contribute to the onset of a multi-faceted response. One of the most important changes caused by waterlogging is the reduction in oxygen availability (hypoxia) to roots, and also to shoots in the case of flooding (Bailey-Serres *et al*., 2012a; Bailey-Serres *et al*., 2012b; Sasidharan *et al*., 2017). Other parameters that change and may affect plants negatively upon waterlogging and flooding include the release of toxic chemical compounds in the soil (*e.g.,* iron) (Setter and Waters, 2003), reduced availability of important nutrients such as nitrates (Setter and Waters, 2003), changes in the soil microbiome (Hartman and Tringe, 2019; Wang *et al*., 2017), or a decrease in light availability because of flood water turbidity.

In recent years, the study of waterlogging stress and of the resulting hypoxic stress has led to the discovery of essential and conserved oxygen-sensing mechanisms in plants (reviewed in (Doorly and Graciet, 2021; Hammarlund *et al*., 2020; Holdsworth and Gibbs, 2020)). An important oxygen-sensing pathway in plants requires a set of oxygen-dependent PLANT CYSTEINE OXIDASE (PCO) enzymes that oxidize the N-terminus of proteins starting with a cysteine residue (Weits *et al*., 2014; White *et al*., 2018; White *et al*., 2017), including a set of group VII ETHYLENE RESPONSE FACTOR (ERF-VII) transcription factors (White *et al*., 2018; White *et al*., 2017) that act as master regulators of the hypoxia response program (reviewed in (Giuntoli and Perata, 2018)). Under normal oxygen conditions, these ERF-VII transcription factors are targeted for degradation by the ubiquitin-dependent N-degron pathway, following oxidation of their N-terminal cysteine residue by PCO enzymes (Gibbs *et al*., 2011; Licausi *et al*., 2011; Weits *et al*., 2014). In contrast, under hypoxic conditions, the activity of PCOs is inhibited, thus preventing the oxidation of ERF-VIIs’ N-terminal cysteine and their subsequent N-degron-mediated degradation. As a result, under hypoxic conditions, ERF-VIIs accumulate in the nucleus, where they bind to *cis*-regulatory elements to activate the hypoxia response program. Notably, the promoters of conserved hypoxia response gene families are not only enriched for *cis*-regulatory elements bound by transcription factors of the ERF-VII family (*e.g.*, the HRPE motif (Gasch *et al*., 2016)), but also for motifs bound by basic helix-loop-helix (bHLH), MYB and WRKY transcription factors (Reynoso *et al*., 2019), thus suggesting the involvement of other families of transcription factors in the regulation of the hypoxia response program (Lee and Bailey-Serres, 2021). This is in agreement with the up-regulation of transcription factor-coding genes belonging to the different families mentioned above as part of hypoxia response in plants (Mustroph *et al*., 2010). Cross-comparisons of transcriptomic datasets from different plant species further allowed the identification of core genes of the hypoxia response program (Mustroph *et al*., 2010; Mustroph *et al*., 2009; Reynoso *et al*., 2019). These include genes involved in (i) the regulation of carbon metabolism in order to facilitate ATP production *via* glycolysis and NAD+ regeneration through fermentation pathways; (ii) the regulation of important signaling pathways (*e.g*., MAPK signaling) and molecules (*e.g*., reactive oxygen species (ROS)) to promote plant tolerance and survival upon hypoxia.

Numerous transcriptomic analyses using either flooding or waterlogging treatments have been conducted with the model plant *Arabidopsis thaliana* (see list in (Brazel and Graciet, 2023)). These studies revealed that core aspects of the transcriptional reprogramming upon waterlogging or flooding are shared with the response to hypoxia (Lee *et al*., 2011; van Veen *et al*., 2016). Furthermore, the transcriptomic comparison of 8 different natural accessions of Arabidopsis, which had been previously identified as being either sensitive or tolerant to flooding (Vashisht *et al*., 2011), indicated that the transcriptional response of roots and shoots differ (van Veen *et al*., 2016). This work identified sets of shoot or root-expressed genes that might contribute to the flooding tolerance phenotype of some natural Arabidopsis accessions (van Veen *et al*., 2016), which could be relevant to improve crop tolerance to waterlogging/flooding. Notably, the transcriptional reprogramming in response to hypoxia and flooding is also accompanied by other genome-wide regulatory mechanisms such as epigenetic changes, chromatin remodelling (Reynoso *et al*., 2019), and changes in translation (Lee and Bailey-Serres, 2019; Reynoso *et al*., 2019).

Waterlogging is an important source of crop losses, however, not all crops are equally affected (de San Celedonio *et al*., 2014; Kaur *et al*., 2020). Barley is particularly sensitive to waterlogging with reported crop losses of up to 20-25% (Byrne *et al*., 2022; de San Celedonio *et al*., 2016; de San Celedonio *et al*., 2014; Liu *et al*., 2020; Miricescu *et al*., 2021; Setter and Waters, 2003). These losses are particularly severe if waterlogging occurs at the heading stage (de San Celedonio *et al*., 2016; de San Celedonio *et al*., 2014; Liu *et al*., 2020; Setter and Waters, 2003), but also at early developmental stages (de San Celedonio *et al*., 2016; de San Celedonio *et al*., 2014). The multifaceted nature of waterlogging stress, as well as the complexity of plant responses to this stress, have made it difficult to identify marker genes of waterlogging tolerance, as well as target genes that could be modified to improve crop tolerance to waterlogging. Nevertheless, in recent years, genetic approaches, as well as linkage mapping and genome-wide association studies (GWAS), have led to the identification of potential QTLs and target genes to improve barley tolerance to waterlogging (Bertholdsson *et al*., 2015; Borrego-Benjumea *et al*., 2020; Broughton *et al*., 2015; Gill *et al*., 2019; Li *et al*., 2008; Manik *et al*., 2022; Zhang *et al*., 2017; Zhang *et al*., 2016; Zhou, 2011). In agreement with the complex response to waterlogging, these QTLs have been identified based on a wide range of phenotypic traits, including leaf yellowing, chlorophyll fluorescence, yield traits, adventitious root formation, aerenchyma formation and ROS levels. QTLs relating to root traits, including the ability to form adventitious roots and aerenchyma under waterlogged conditions (Manik *et al*., 2022; Zhang *et al*., 2016), may be of particular relevance considering their known roles in facilitating water and nutrient uptake as well as in oxygen distribution during waterlogging (Zhang *et al*., 2015).

Despite the recent progress made in the identification of QTLs for waterlogging tolerance and the realization that transcriptomics may be used to identify potential candidate genes relevant to waterlogging tolerance (Lee *et al*., 2011; Reynoso *et al*., 2019; van Veen *et al*., 2016), only few such studies have been conducted with barley. Recent studies have characterized the transcriptomic response of waterlogging-tolerant and sensitive varieties (Borrego-Benjumea *et al*., 2020; Luan *et al*., 2022). From these datasets, the authors identified a handful of genes that could be of importance for waterlogging tolerance mechanisms in barley (Luan *et al*., 2022). In another study, a comparative proteomic analysis of one waterlogging-sensitive and one waterlogging-tolerant variety identified proteins that accumulate differently in waterlogging sensitive or tolerant germplasms (Luan *et al*., 2018a; Luan *et al*., 2018b).

Here, to dissect the transcriptomic response of barley to waterlogging, we selected two 2-row and two 6-row winter barley varieties based on the differential expression of known hypoxia response marker genes after waterlogging treatment. These varieties were chosen from the Association Genetics Of UK Elite Barley (AGOUEB) population (Thomas *et al*., 2014) and from the list of recommended barley varieties in Ireland (where the study was conducted; list at the time the experiments were carried out). Our transcriptomic study focused on roots because of their central role in mediating waterlogging tolerance (Zhang *et al*., 2015). We also included the model spring barley variety *Golden Promise* to provide a reference dataset to the community (this variety is widely used to target specific genes using molecular approaches). We identified sets of common genes that are consistently differentially expressed in barley in response to waterlogging. Hierarchical clustering identified groups of genes with intrinsic differential expression between varieties prior to waterlogging stress. Low or elevated expression of some of these genes could ‘predispose’ some varieties to waterlogging tolerance or sensitivity. In sum, the datasets presented serve as an additional reference for the study of waterlogging response in barley and provide insights into potential avenues of research to improve waterlogging tolerance in this crop.

## Materials and Methods

### Plant material

Cultivars used in this study (Supp. Table S1) included winter varieties that originated from the ‘Association Genetics of UK Elite Barley’ (AGOUEB) population of barley (Thomas *et al*., 2014), as well as the model spring variety *Golden Promise* (obtained from Teagasc, Oak Park, Ireland) and a winter variety, *Infinity* (obtained from Teagasc, Oak Park, Ireland), which was on the recommended list of barley varieties in Ireland (where the study was conducted) at the time of the initial field trial whose results were taken into account for varietal selection (Byrne *et al*., 2022). Selected varieties for further characterization, including transcriptome profiling, were *Golden Promise* (spring variety, 2-row), *Regina* and *Infinity* (winter varieties, 2-row), *Passport* and *Pilastro* (winter varieties, 6-row).

### Plant growth conditions

Plants were grown in a plant growth room under long-day conditions (16 h light/8 h dark) at 15°C (constant temperature), approximately 45% relative humidity. Light intensity was determined to be ∼138 µmol/m^2^/s and was provided by LED bulbs (Philipps LED tubes High Output, T8 20W/865).

### Soil preparation and seed germination

Commercial John Innes N°2 (Vitax; UK) soil was soaked in water after filling round pots of 9-cm diameter and 9-cm height without compressing the soil. Untreated seeds were sown directly in soil at a depth of 2 cm. The sown seeds were stratified in the dark for 14 d at 4°C to ensure homogenous germination. After cold treatment, the pots were transferred to the plant room for germination and growth.

### Waterlogging

Plants were grown as indicated above for ten days (corresponding to L1/L2 stage) for transcriptomic experiments, and for 10-14 d (corresponding to L1/L2 stage) for phenotypic characterization. The pots were then transferred to a large tub, which was subsequently filled with tap water up to 1 cm above soil level. The water level was kept constant throughout the duration of the experiment (Figure 1A). The plants were kept in the same growth conditions and were treated for 15 d for phenotypic characterization, or for shorter periods of time, as indicated in the text. Control plants were left in the same growth conditions but received normal watering (every two days, avoiding any standing water in the trays). For the recovery period, plants were taken out of the water and kept in the same growth conditions with a normal watering regime.

**Figure 1:**
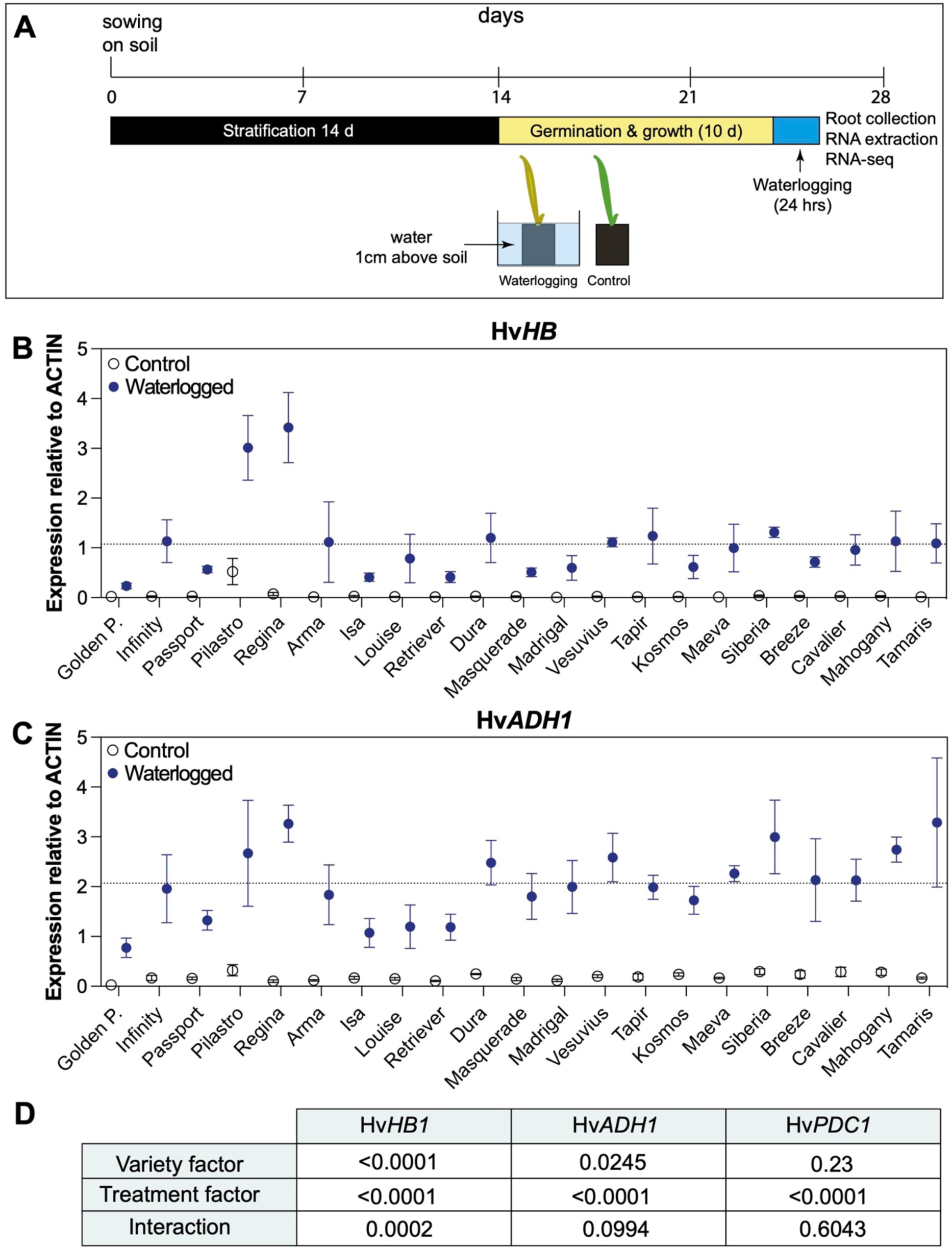
Differential expression of selected hypoxia response genes in 21 different barley varieties. (A) Experimental design of waterlogging experiments (see Materials and Methods for details). (B) Hv*HB* expression relative to *ACTIN* in untreated (open symbols; normal watering) and waterlogged (blue symbols) plants after 24 h of treatment. The dashed line corresponds to the average relative expression of Hv*HB* for waterlogged samples of all varieties and biological replicates. (C) Hv*ADH1* expression relative to *ACTIN* in untreated (open symbols) and waterlogged (blue symbols) plants after 24 h of treatment. The dashed line corresponds to the average relative expression of Hv*ADH1* for waterlogged samples of all varieties and biological replicates. (D) *p*-value results of two-way ANOVA analysis of the data presented in (B) and (C), and in Supp. Figure S2. Data shown in (B) and (C) are from three biological replicates, except for *Golden Promise* for which 4 biological replicates were used. Errors bars indicate SEM. Each biological replicate was obtained by pooling the root systems from 3 plants prior to grinding for RNA extraction. Note: expression values for *Infinity* shown in (B) and (C) were published in (Miricescu *et al*., 2021).

### Height measurements

Plant height was taken from the soil surface to the tip of the tallest leaf at the indicated time points.

### Total RNA extraction

Roots of plants grown under control conditions or subjected to waterlogging for were rinsed under running tap water, and frozen in liquid nitrogen. The tissue was ground in liquid nitrogen and total RNA was extracted using Spectrum^™^ Plant Total RNA kit (Sigma-Aldrich). For each condition (waterlogged or untreated), the root systems of 3 plants of the same variety were pooled prior to grinding for total RNA extraction. This experiment was conducted independently at least 3 times to obtain samples from at least 3 biological replicates, as indicated in the figure legends.

### Reverse transcription coupled to quantitative PCR (RT-qPCR)

Total RNA was reverse transcribed using an oligo(dT)_18_ primer and the RevertAid reverse transcriptase (Thermo) according to the manufacturer’s instructions. For reverse transcription reactions, 1 µg of total RNA was used. cDNA obtained was used for qPCR with a Lightcycler 480 (Roche). Each PCR reaction mix contained 5 μL of 2x SYBR green master 1 (Roche), 1 μL cDNA, 1 μL of 10 μM primers and 3 μL of molecular biology grade water.

LightCycler melting curves were obtained to check for single peak melting curves for all amplification products. The second derivative maximum method was used to analyze the amplification data. The resulting Cp values were converted into relative expression values using the comparative Ct method (Livak and Schmittgen, 2001). All primer sequences are provided in Supp. Table S2. One reference gene (Hv*ACTIN*) was used to normalize the RT-qPCR data following the screening of a set of reference genes during waterlogging (Supp. Fig. S1).

### Next-generation sequencing of RNA samples

For RNA-seq analysis, waterlogging treatment was applied as outlined above for 24 h. For each condition (waterlogged or untreated), the root systems of 3 plants of the same variety were pooled prior to grinding for total RNA extraction. This experiment was conducted independently 3 times to obtain samples from 3 biological replicates (i.e. for each variety, 6 RNA samples were sent for sequencing, corresponding to 3 biological replicates for the untreated plants and 3 biological replicates for the waterlogged plants). Following RNA extraction, RNA integrity was assessed using an Agilent 2100 Bioanalyzer (Agilent). All RNA samples had RIN values > 7.0. Library preparation and single ended 50 bp next-generation sequencing was performed by BGI Genomics (Hong Kong) using the DNBseq sequencing platform.

### Analysis of RNA-seq data

The third release (Morex V3) of the Morex barley genome was downloaded from e!DAL - Plant Genomics and Phenomics Research Data Repository (PGP) (https://edal-pgp.ipk-gatersleben.de/) (Mascher, 2021). Raw RNA-sequencing reads were aligned to Morex V3 using *bowtie2* (v2.4.5) (Langmead and Salzberg, 2012). Files were converted from .sam to .bam files and indexed using *samtools* (v1.15.1) (Danecek *et al*., 2021). Gene abundance was estimated using *stringtie* (v2.1.7) (Pertea *et al*., 2015) (Supp. Table S3). Differential gene expression analysis was performed using the Bioconductor package *DeSeq2* (Love *et al*., 2014) in *R* (Team, 2022) using a design in which the factors Variety and Treatment were combined into a single factor to model multiple condition effects. The results of multiple comparisons were extracted and filtered by adjusted *p*-value < 0.05 (Supp. Table S4). Principal component analysis (PCA) was performed using *pcaExplorer* (v2.20.2) (Marini and Binder, 2019) in *RStudio* (v2022.02.0+443). Gene ontology (GO) analysis was performed using *ShinyGO* (v0.75c) (Ge *et al*., 2020). Read count values were transformed by variance stabilising transformations (VST) to normalise for library size and composition (Supp. Table S5). Venn diagrams were generated using InteractiVenn (Heberle *et al*., 2015).

The means of VST read counts from 3 biological replicates generated from *DeSeq2* were filtered for *k*-means clustering as follows. *Deseq2* DEG analysis was performed and results for the 25 comparisons shown in Supp. Fig S6A were extracted. A list of all 11,613 DEGs filtered by adjusted *p*-value < 0.05 was generated by combining DEGs from waterlogged to control samples from the same variety, DEGs from each control to every other control sample and DEGs from each waterlogged to every other waterlogged sample. Next, the DEGs with a mean of <10 normalised reads were removed leaving 10,882 genes for clustering analysis. Clustering analysis was performed using the *k*-means function in *R* (v3.6.2) with the arguments centers=25, nstart = 1000, iter.max = 300 and algorithm = “Lloyd” (gene cluster assignment annotated in (Supp. Table S4)). Clustering heatmaps were generated using *ComplexHeatmap* (v2.10.0) (Gu *et al*., 2016) in *R*. Plots were generated using *ggplot2* (v3.3.6) (Wickam, 2016) and modified for style in Adobe Illustrator 2022.

To compare the datasets obtained to a previously published RNA-seq dataset, raw sets were downloaded from NCBI’s Gene Expression Omnibus (GSE144077) (Borrego-Benjumea *et al*., 2020) and Sequence Read Archive (PRJNA889532) (Luan *et al*., 2022). Raw RNA-sequencing reads were aligned to Morex V3 as described above. Differential gene expression analysis of waterlogging vs control for each variety was performed on 0 h control and 24 h waterlogging datasets for the varieties Franklin and TX9425 from Luan *et al*. (2022) as described above. The same differential gene expression analysis was also performed on 72 h control and 72 h waterlogging datasets for the varieties Deder 2 and Yerong from Borrego-Benjumea *et al*. (2020).

To compare the datasets to a previously published list of 28 genes found in QTLs for waterlogging tolerance, we downloaded the gene list from Supp. Table 3 in (Manik *et al*., 2022). This gene list used Morex V1 (r1) gene IDs. To compare this list to our data, we performed a BLASTN search of the CDS of each of these genes to the Morex V3 genome using *Phytozome 13* (Goodstein *et al*., 2012) and used the transcript with the highest percentage identity to a Morex V3 transcript for the comparison (Supp. Table S4).

## Results

### Characterization of the transcriptional response of different barley varieties using hypoxia response marker genes

The previous characterization of 403 varieties from the AGOUEB collection under waterlogged and control conditions in the field (Byrne *et al*., 2022) provided some information on the differential physiological response of these varieties to waterlogging, while also highlighting the difficulties associated with the study of waterlogging tolerance under field conditions. Based on this initial study, we selected a subset of 20 varieties that behaved differently under waterlogged conditions, with the aim to assess their transcriptional response to waterlogging under controlled conditions. The variety *Golden Promise* was also included as a model (spring) barley variety that is widely used to generate targeted mutations through *Agrobacterium*-mediated transformation. After identifying Hv*ACTIN* as a suitable reference gene in waterlogged roots (Supp. Figure S1A), we first carried out a time course experiment with four selected varieties and determined the relative expression of *HEMOGLOBIN1 (HB)*, *ALCOHOL DEHYDROGENASE 1 (ADH1)* and *PYRUVATE DECARBOXYLASE 1 (PDC1)*. These genes are hypoxia response markers commonly used to monitor waterlogging response (Loreti *et al*., 2020; Mendiondo *et al*., 2016; Mustroph *et al*., 2010). This initial analysis indicated that (i) the expression of these three hypoxia response genes peaked around 24 h after the onset of the waterlogging treatment; and (ii) there were differences between the four varieties in terms of the amplitude of the transcriptional up-regulation (Supp. Figure S1B). For example, at 24 h after the beginning of the waterlogging treatment, Hv*HB* was expressed at higher levels in *Pilastro* compared to *Arma*, *Tapir* and *Masquerade*. In addition, the up-regulation of Hv*HB* and Hv*ADH1* was stronger than that of Hv*PDC1*, possibly making these first two genes more suitable to identify varieties with differential transcriptional regulation of the hypoxia response program.

We next tested the expression of Hv*HB*, Hv*ADH1* and Hv*PDC1* at 24 h of waterlogging in the set of 21 barley varieties we selected based on (Byrne *et al*., 2022) (Figure 1A). As expected, under normal watering conditions, most varieties had low levels of expression for each of the hypoxia response genes, with the exception of *Pilastro*, which exhibited higher Hv*HB* expression (Figures 1B, 1C and Supp. Figure S2). Waterlogging stress triggered the up-regulation of the hypoxia response genes in all varieties tested, but some differences in their response were observed. As previously, differences were more marked with Hv*HB* and Hv*ADH1* (Figure 1B-D) than with Hv*PDC1* (Supp. Figure S2), making these first two genes more suitable to identify varieties with a differential transcriptional response to waterlogging. After 24 h of waterlogging, some varieties reached higher expression levels than the average relative expression observed for these genes in the population of 21 varieties. These included *Pilastro* and *Regina* for both Hv*HB* and Hv*ADH1*, as well as *Dura*, *Vesuvius*, *Siberia*, *Mahogany* and *Tamaris* for Hv*ADH1*. In contrast, some varieties showed reduced up-regulation of hypoxia response genes compared to the population average. For example, varieties such as *Golden Promise*, *Passport*, *Isa*, *Louise* and *Retriever* had lower expression levels of both Hv*HB* and Hv*ADH1* compared to the average of the population. Other varieties, such as *Infinity*, had average expression levels of Hv*HB* and Hv*ADH1*.

Based on these results, *Pilastro* and *Regina* were chosen as representative 6-row and 2-row varieties, respectively, that potentially exhibit a stronger transcriptional response to waterlogging, while *Passport* was selected as a 6-row variety that had a more dampened transcriptional response. We also included *Infinity* as a 2-row variety for further characterization because it is on the recommended list in Ireland (where this study was carried out) and has an average transcriptional response to waterlogging. As highlighted above, *Golden Promis*e was included as a reference variety.

### Effect of waterlogging on the growth of selected varieties

The growth of the five selected varieties was characterized under controlled conditions following a 2-week waterlogging treatment, and a 6-week recovery period with normal watering. Height measurement showed that the growth of *Infinity* and *Passport* was more affected by waterlogging than that of *Pilastro*, *Regina* and *Golden Promise*, whose height was mostly unaffected by the treatment (Figure 2A). In addition, *Golden Promise*, *Infinity* and *Pilastro* produced fewer tillers, whereas *Regina*’s tiller number was largely unaffected (Figure 2B). Because root traits have been shown to be important for waterlogging tolerance (Zhang *et al*., 2015), we determined the length of the primary root, as well as the number of seminal roots after 2 weeks of waterlogging. The length of the primary root was reduced upon waterlogging stress for all varieties except *Regina* (Figure 2C), while the number of seminal roots increased in all varieties and the difference was statistically significant for all except *Infinity* (Figure 2D). This phenotypic characterization suggests that a variety such as *Regina*, which has a stronger transcriptional response for Hv*ADH1* and Hv*HB*, appears to be less impacted by waterlogging than other varieties.

**Figure 2:**
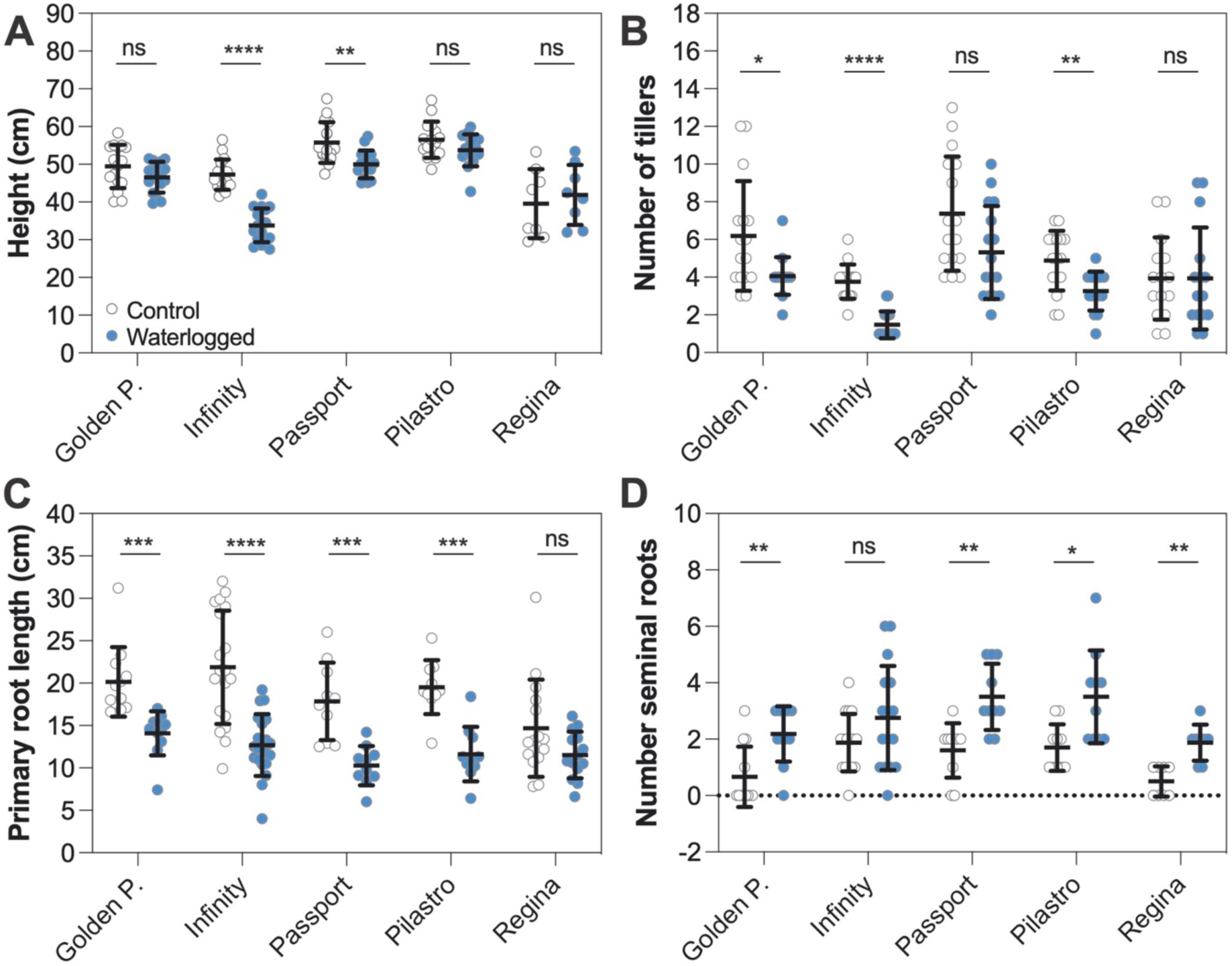
Effect of waterlogging on shoot and root growth. (A) Plant height after 14 d of waterlogging and a 6-week recovery period. For *Golden Promise*, *Passport* and *Pilastro*, three biological replicates (4 plants per biological replicate) were carried out. For *Regina* and *Infinity*, two biological replicates (4 plants per replicate) and 3 biological replicates (4 to 9 plants per replicate) were carried out, respectively. (B) Number of tillers after 14 d of waterlogging and a 6-week recovery period. Three biological replicates were carried out with at least 4 plants per replicate. (C) Length of the primary root after 14 d of waterlogging. For *Golden Promise*, *Passport*, *Pilastro* and *Regina*, three biological replicates with 2 to 7 plants each were carried out. For *Infinity*, 5 biological replicates were carried out with 3 to 4 plants per replicate. (D) Number of seminal roots after 14 d of waterlogging. For *Golden Promise*, *Passport*, and *Pilastro*, three biological replicates (2 to 4 plants per replicate) were carried out. For *Regina*, 2 biological replicates were conducted (4 plants per replicate). For *Infinity*, 4 biological replicates were carried out (4 plants per replicate). In (A-D), data from different biological replicates is color coded. Error bars correspond to standard deviations. Statistical analysis: multiple unpaired t-test with multiple comparison correction (Holm-Sidak method). Asterisks: * p≤ 0.05; ** p≤0.01; *** p≤0.001; **** p≤0.0001; ns: not statistically significant.

### Overview of transcriptional responses to waterlogging in root tissue of selected varieties

To determine the transcriptional responses of *Pilastro*, *Regina*, *Passport*, *Infinity* and *Golden Promise* to waterlogging, we subjected these varieties to waterlogging for 24 h. This time point was chosen based on the time course experiment described above that showed peak expression of Hv*HB*, Hv*ADH1* and Hv*PDC1* at 24 h of waterlogging. Total RNA was extracted from roots for RNA-seq analysis. Reads obtained were aligned to the Morex barley genome (version 3; Supp. Table S3 and Supp. Figure S3A-C) and differential gene expression (adjusted *p-*value < 0.05) was determined for each of the waterlogged varieties relative to the corresponding untreated variety (Supp. Table S4). A Principal Component Analysis (PCA) indicated that the samples separated based on (i) the treatment (PC1: 35.82% variance) (Figure 3A); (ii) whether they were winter or spring varieties (PC2: 15.64% variance; Supp. Figure S3D); or (iii) whether they were 2-row or 6-row varieties (PC3: 11.34% variance; Supp. Figure S3E). In this analysis, the varieties *Golden Promise*, *Infinity* and *Passport* could also be separated from *Pilastro* and *Regina* (Figure S3F; PC4: 9.92% variance). This suggests that there are underlying differences between these two groups of varieties that are not explained by a known variable.

**Figure 3:**
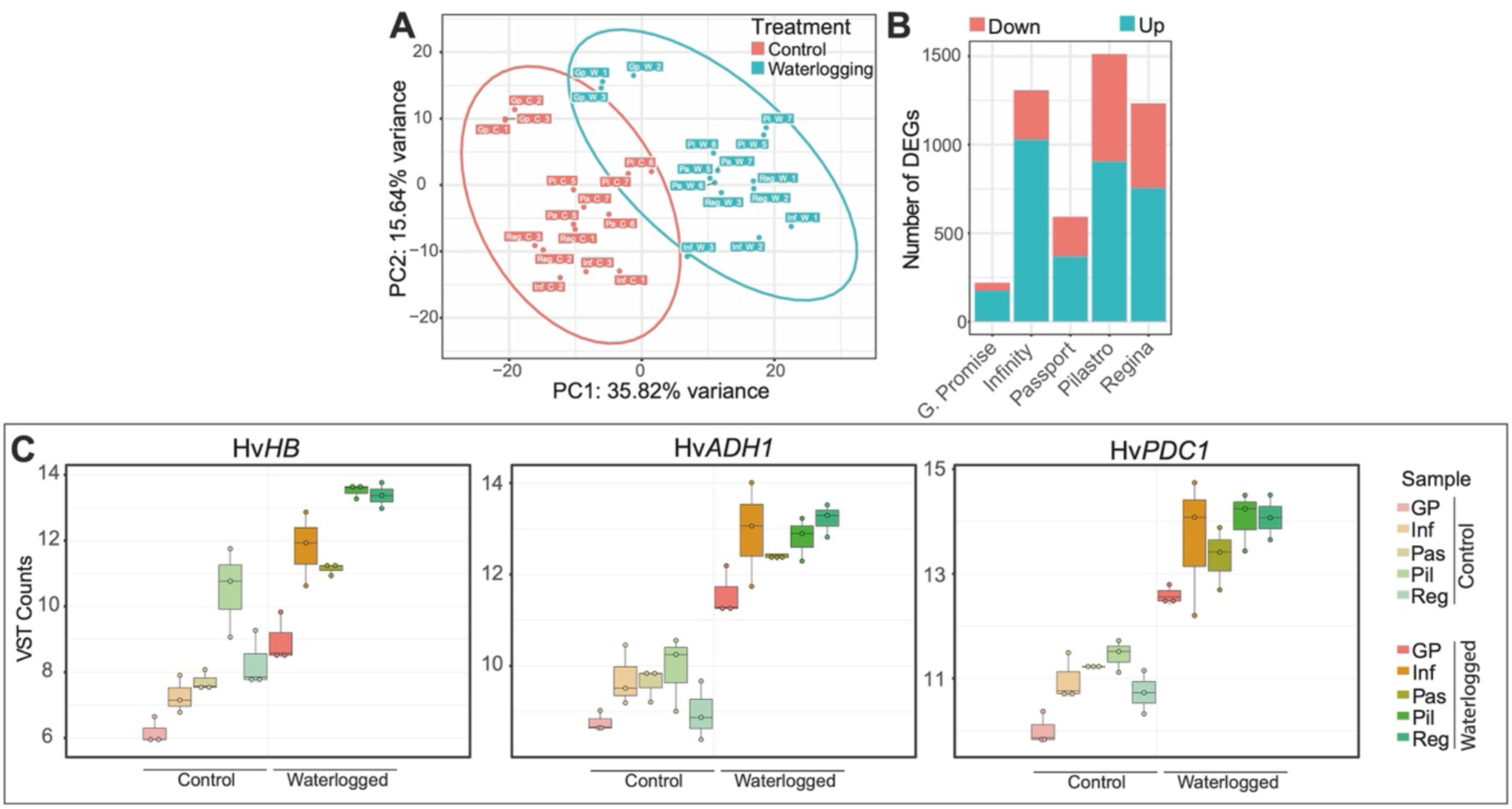
Results of RNA-seq experiments. (A) PCA plot showing Principal Component 2 (PC2) *versus* PC1 with ellipses showing sample grouping by treatment. (B) Absolute number of DEGs (filtered by adjusted *p*-value<0.05) in waterlogged *versus* control samples for each variety. (C) Box plots showing variance stabilised transformed (VST) read counts for Hv*HB* (HORVU.MOREX.r3.4HG0394960), Hv*ADH1* (HORVU.MOREX.r3.4HG0345760.1) and Hv*PDC1* (HORVU.MOREX.r3.4HG0384560) in control and waterlogged samples (*Golden Promise* (GP), *Infinity* (Inf), *Passport* (Pas), *Pilastro* (Pil) and *Regina* (Reg)).

The number of differentially expressed genes (DEGs) identified in the different varieties varied considerably: for *Golden Promise* and *Passport* we identified 220 and 592 DEGs (respectively), while *Infinity*, *Pilastro* and *Regina* had a substantially higher number of DEGs (1,307; 1,512; and 1,233, respectively). In all varieties tested, the proportion of up-regulated genes was higher than that of down-regulated genes (Figure 3B). We verified in our datasets the expression of the Hv*HB*, Hv*ADH1* and Hv*PDC1* genes whose up-regulation was initially used to monitor the transcriptional response of the different varieties to waterlogging by RT-qPCR (Figure 1). Induction of all three genes upon waterlogging was found (Figure 3C). We further identified additional homologs of Hv*ADH1* and Hv*HB* and confirmed that these homologs were also up-regulated in response to waterlogging (Supp. Figure S4).

### Genome-wide transcriptional reprogramming of barley in response to waterlogging

Our RNA-seq experiment identified 2,865 genes that were differentially expressed in at least one variety under waterlogged conditions compared to control plants (Supp. Table S4). The variety with the highest proportion of unique DEGs was *Pilastro* (>50% unique DEGs), while the majority of DEGs identified for *Golden Promise* and *Passport* were shared with other varieties (Figure 4A) (<25% unique DEGs in these 2 varieties). Among the 2,865 DEGs, 1,078 genes were called as differentially expressed in more than one variety. Except for 7 genes, the directionality of gene expression change was the same in all varieties, thus further highlighting the similarities in the response of these varieties to waterlogging. As expected based on the transcriptional response of Arabidopsis to waterlogging, Gene Ontology (GO) analysis of the 1,811 DEGs that were up-regulated in at least one variety identified over-represented terms related to hormone and stress responses, oxidoreductase activity, kinase activity, as well as carbohydrate metabolic processes (Figure 4B and D). GO analysis of the 1,061 DEGs that were down-regulated in response to waterlogging identified terms related to lipid and carbohydrate metabolism, transmembrane transporter activities and cell periphery, as well as oxidoreductase activity and proteolysis (Figure 4C and E). All of these terms are consistent with functional GO categories known to be central to the transcriptional response of plants to waterlogging.

**Figure 4:**
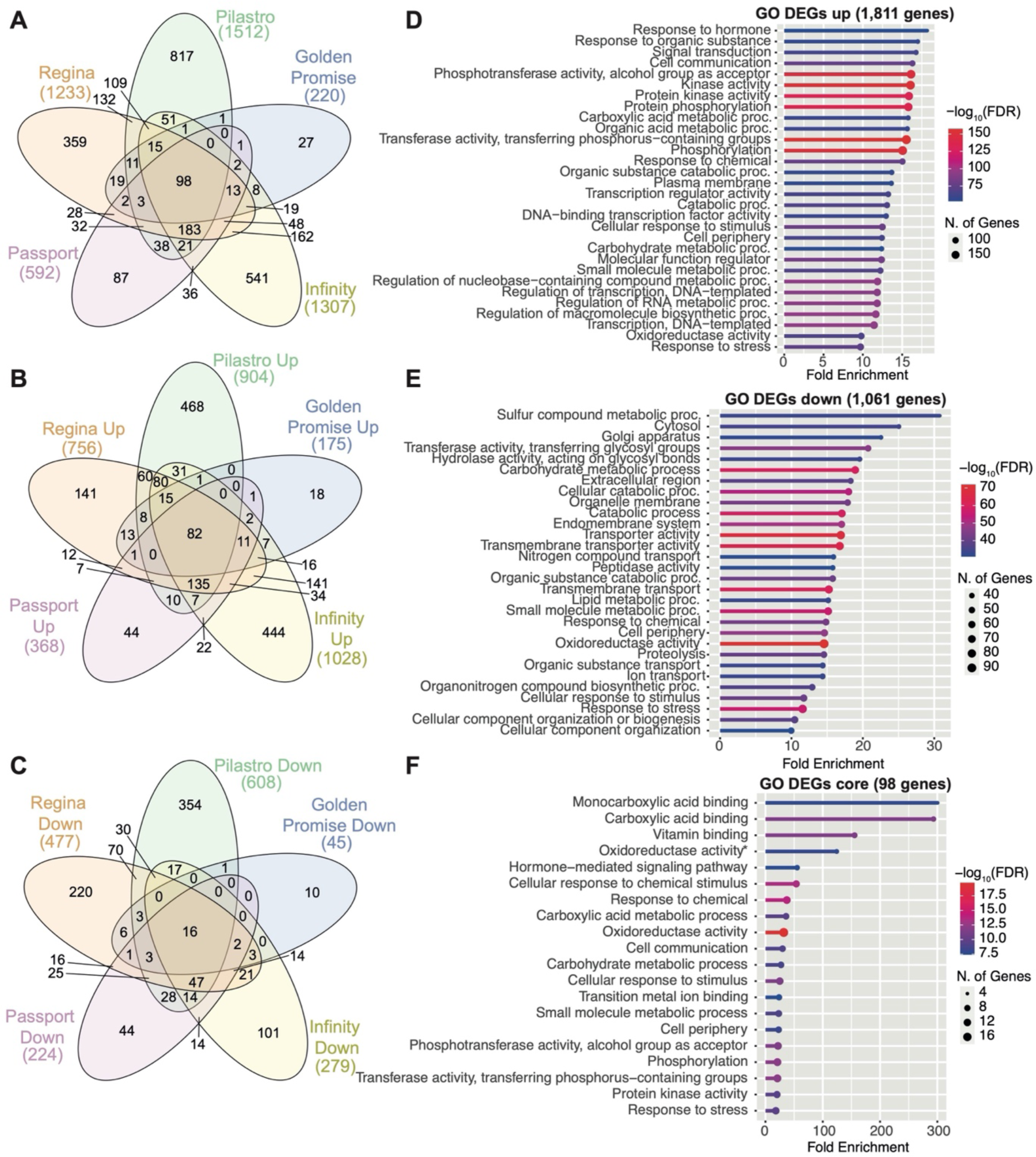
Comparison of *waterlogging* response genes in barley. (A) Overlap between the DEGs identified in the different varieties. (B) Overlap analysis of genes up-regulated in waterlogged samples compared to control samples for each variety. (C) Overlap between datasets for genes that are down-regulated in waterlogged samples compared to the controls in each variety. (D) The 30 most highly enriched GO terms among the 1,811 DEGs that are up-regulated in at least one variety in response to waterlogging. (E) The 30 most highly enriched GO terms enriched among the 1,061 DEGs that are down-regulated in any variety in response to waterlogging. (F) The 20 most enriched GO terms among the 98 DEGs common to all varieties when comparing waterlogged to control samples for each variety.

A total of 98 genes were detected as differentially expressed in the five varieties (Figure 4A), and their directionality of expression change was the same across all datasets. The majority of these genes were up-regulated in response to waterlogging (82 genes; Figure 4B and Supp. Table S4) and included homologs of known core hypoxia response genes in Arabidopsis (Mustroph *et al*., 2009) such as *ADH1* (HORVU.MOREX.r3.4HG0345760, HORVU.MOREX.r3.1HG0082960 and HORVU.MOREX.r3.4HG0345740), *PDC1* (HORVU.MOREX.r3.4HG0384560), *HB1* (HORVU.MOREX.r3.4HG0394960, HORVU.MOREX.r3.7HG0715010) and a respiratory burst oxidase (*RBOH*) (HORVU.MOREX.r3.5HG0444960). Furthermore, this set included homologs of ERF-VII transcription factors, such as *HRE1* (HORVU.MOREX.r3.4HG0405650 and HORVU.MOREX.r3.4HG0405650), *HRE2* (HORVU.MOREX.r3.5HG0497970) and *RAP2.3*, (HORVU.MOREX.r3.1HG0060770). This result is in agreement with the central role of ERF-VII transcription factors as master regulators of hypoxia and waterlogging response. Sixteen DEGs common to all datasets were down-regulated in response to waterlogging (Figure 4C), including three peroxidase genes (HORVU.MOREX.r3.2HG0215440, HORVU.MOREX.r3.2HG0119650, HORVU.MOREX.r3.2HG0119630) a nitrate transporter (HORVU.MOREX.r3.7HG0700030), and an aldehyde dehydrogenase (HORVU.MOREX.r3.2HG0173750). GO analysis of the set of 98 common DEGs (Figure 4F) revealed largely similar terms to those identified for all up-regulated DEGs (Figure 4D), including GO terms such as oxidoreductase activity, carbohydrate metabolic process, and hormone-mediated signaling.

We next compared our RNA-seq results to two recently published RNA-seq datasets. Luan et al. analyzed the transcriptomic response of 4-leaf stage Franklin and TX9425 barley roots to waterlogging for 24 or 72 h (Luan *et al*., 2022). These were two-row spring varieties, with Franklin showing sensitivity and TX9425 showing tolerance to waterlogging (Luan *et al*., 2022). In addition, Borrego-Benjumea *et al*. analyzed the transcriptomic response of waterlogged barley roots in 2-week-old *Yerong* and *Deder2* for 72 or 120 h (Borrego-Benjumea *et al*., 2020). Both varieties had been chosen because of their tolerance to waterlogging (Li *et al*., 2008; Takeda, 1989). Raw RNA-seq data were analyzed using the same pipeline as for our datasets. PCA analysis of all datasets showed that most of the variation was due to the origin of the dataset (Figure S5A; PC1: 68.39% variance), likely because waterlogging stress is known to vary considerably based on experimental differences, such as soil type. As expected, the different samples could also be separated based on treatment (Figure S5A; PC2: 17.03% variance). To facilitate the comparison between the different datasets, we selected the most similar timepoints to those of our study (*i.e*., 0 h control and 24 h waterlogged for Luan *et al*.; and 72 h control and waterlogged for Borrego-Benjumea *et al*.). Despite the differences in barley varieties, age of the plants and duration of the waterlogging treatment, a statistically significant overlap was observed between both the up and down-regulated genes in our datasets and in those recently published datasets DEGs (Supp. Figure S5B and S5C) (Borrego-Benjumea *et al*., 2020; Luan *et al*., 2022). GO analysis of the 448 DEGs that were up-regulated in all datasets identified terms related to metabolism and oxidoreductase activity (Supp. Figure S5F), again in agreement with known key aspects of plant responses to waterlogging. Genes associated with transporter activity were found among the 309 shared down-regulated genes (Supp. Figure S5G).

### Identification of unique expression signatures between varieties

To compare the responses to waterlogging between the 5 varieties used in this study, *k*-means clustering was performed on 10,882 genes identified as differentially expressed in: (i) at least one variety when comparing waterlogged to control samples for each variety, or (ii) between the control samples (*i.e*., these have intrinsic differential expression between at least 2 varieties in the absence of waterlogging), or (iii) between the waterlogged samples from different varieties (*i.e*., genes that are differentially expressed between at least 2 varieties under waterlogging) (Supp. Figure S6A). Clustering analysis revealed distinct gene expression patterns between samples (Figure 5A). A number of clusters contained genes that were up-regulated in multiple varieties in response to waterlogging compared to untreated plants (*i.e.*, clusters 4, 7, 8, 14, and 21). The largest cluster (#14) comprised 802 genes (Supp. Table S4) with a centroid expression that was higher in all varieties in response to waterlogging (Figure 5B and Supp. Figure S6). Cluster 14 included homologs of known hypoxia response genes such as those described above in the common set of 82 up- regulated DEGs in barley (Figure 4B), as well as homologs of Arabidopsis *RAP2.12* (HORVU.MOREX.r3.5HG0481240), additional homologs of *RAP2.3*, (HORVU.MOREX.r3.2HG0112960), and homologs of *ADH1* (HORVU.MOREX.r3.2HG0115170, HORVU.MOREX.r3.3HG0252910) (Supp. Figure S4A and B). Furthermore, transcription factors (TFs) from the ERF, bHLH, bZIP (basic leucine zipper), NAC (NAM, ATAF1,2, CUC2) and WRKY families were also identified in cluster 14 (Supp. Table S4), all of which belong to families with established and conserved roles in the regulation of hypoxia response genes (Lee and Bailey-Serres, 2021; Mustroph *et al*., 2010; Reynoso *et al*., 2019). GO analysis of the 802 genes in cluster 14 identified terms related to hormone and stress responses, oxidoreductase and kinase activity and metabolism (Figure 5C).

**Figure 5:**
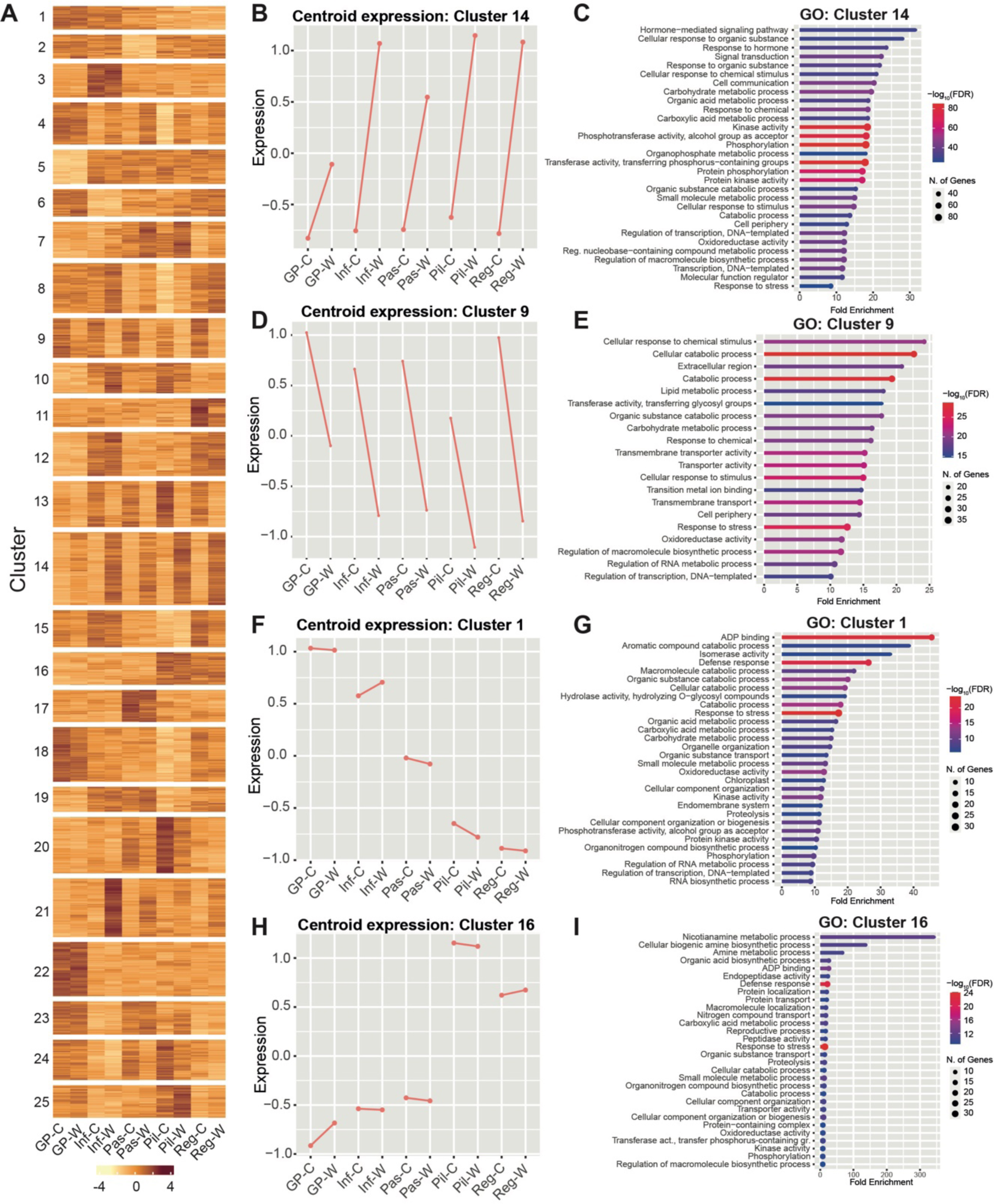
Clustering analysis of DEGs in waterlogged compared to control samples. (A) Heatmap showing the results of *k*-means clustering (*k*=25) of 10,882 DEGs (see Supp. Figure S6A for selection criteria). Image generated using *ComplexHeatmap* in *R*. (B) Plot of centroid expression for genes in cluster 14. (C) The 20 most highly enriched GO terms among the 802 genes identified in cluster 14. (D) Plot of centroid expression for DEGs in cluster 9. (E) The 20 most highly enriched GO terms among the 435 genes identified in cluster 9. (F) Plot of centroid expression for genes in cluster 1. (G) The 20 most highly enriched GO terms among the 257 genes from cluster 1. (H) Plot of centroid expression for genes in cluster 16. (I) The 20 most highly enriched GO terms among the 359 genes from cluster 16.

In contrast, clusters 5, 9, 13, 15, 20 and 24 comprised genes that were down- regulated in response to waterlogging in multiple varieties. Cluster 9 contained 435 genes that were down-regulated in response to waterlogging (Figure 5D and Supp. Figure S6), including 14 of the 16 genes that were down-regulated in waterlogged samples from all varieties (Figure 4C and Supp. Table S4). Among these genes was an aminotransferase (HORVU.MOREX.r3.2HG0190680), whose top-scoring Arabidopsis homolog is γ-aminobutyric acid transaminase (GABA-T; AT3G22200). In Arabidopsis, the expression of this gene is also reduced in response to hypoxia (Branco-Price *et al*., 2008). Another member of cluster 9 whose expression was downregulated in all varieties in response to waterlogging is a cation exchanger (HORVU.MOREX.r3.3HG0266310). In Arabidopsis, the closely related Ca^2+^/proton exchanger 11 (CAX11) is important to maintain cytosolic Ca^2+^ homeostasis and was shown to be down-regulated in response to hypoxia and waterlogging (Wang *et al*., 2016). Multiple transcription factors from the heat-shock and MYB transcription factor families were also found in cluster 9. This is consistent with the observation that differentially expressed MYB transcription factors were predominantly found to be down- regulated in response to waterlogging in *Yerong* and *Deder2* (Borrego-Benjumea *et al*., 2020). GO analysis of the 435 genes in cluster 9 showed an enrichment for terms related to lipid and carbohydrate metabolism, transmembrane transporter activities and cell periphery (Figure 5E), similarly to the GO terms identified in all genes down-regulated in response to waterlogging (Figure 4E).

Clustering analysis also identified gene expression signatures that were specific to individual varieties: cluster 1 (257 genes) contained genes whose expression was highest in the barley varieties *Golden Promise* and *Infinity* and intermediate in *Passport*. In contrast, the expression of these genes was markedly lower in the varieties *Pilastro* and *Regina* (Figure 5F and Figure S6). GO analysis of genes in cluster 1 identified terms related to catabolic and metabolic processes (Figure 5G), which could be relevant to waterlogging tolerance/sensitivity. In contrast, genes in cluster 16 (359 genes) had an opposite behavior, in that they were more highly expressed in *Pilastro* and *Regina*, but were only expressed at low levels in the other varieties (Figure 5H and Figure S6). GO analysis of genes in cluster 16 revealed an enrichment for genes associated with diverse processes found in other clusters or sets of genes, except the term ‘nicotianamine metabolic process’ which was only found in the GO analysis of cluster 16 genes. This GO term relates to metal (including iron) homeostasis, which could be of relevance to waterlogging stress tolerance because of the increased uptake of metals and their toxicity to plants in waterlogged conditions. Notably, genes in clusters 1 and 16 were not differentially expressed in response to waterlogging. Instead, it is their intrinsic (or varietal-specific) expression level that differs between the samples.

## Discussion

Maintaining yields in the context of global climate change is a challenge, as crops are likely to experience a wider range of stresses simultaneously or sequentially. Accordingly, efforts are now made to improve crop resilience to multiple abiotic and biotic stresses, as opposed to tolerance to a specific stress. At the same time, the trade-off between the gain of new traits and yield also requires careful consideration (Bailey-Serres *et al*., 2019). Waterlogging in particular causes important crop losses and will become more prevalent in some regions. Several QTLs have been identified with genes that could be targeted to improve crop tolerance to waterlogging, but the role of candidate genes often needs to be validated, as well as their suitability to improve crop tolerance to waterlogging. Alternatively, targeted mutations in genes coding for key hypoxia response components, such as genes of the N-degron protein degradation pathway, have yielded Arabidopsis and barley lines that are more tolerant to waterlogging and other abiotic stresses (Mendiondo *et al*., 2016; Vicente *et al*., 2017), albeit with a potential trade-off on their ability to resist infection by some pathogens (de Marchi *et al*., 2016; Gravot *et al*., 2016; Vicente *et al*., 2019).

Barley is an essential crop that exhibits varietal differences in its sensitivity to waterlogging. The identification of potential candidate genes that can be targeted or used as markers to identify waterlogging tolerant varieties would be of importance. One approach to identifying candidate targets and markers is the comparison of genome-wide transcriptional responses of different varieties to waterlogging. Here, after identifying a time point (24 h) at which the expression of well-known hypoxia response marker genes peaked, we screened a subset of 20 winter varieties to identify some with a differential transcriptional response to waterlogging. Based on this analysis, we determined the genome-wide transcriptional response to waterlogging of 4 winter barley varieties and of the model spring variety *Golden Promise*. In agreement with previous transcriptional studies with a range of plants and crops (Borrego-Benjumea *et al*., 2020; Lee *et al*., 2011; Lee and Bailey-Serres, 2021; Luan *et al*., 2022; Luan *et al*., 2023; Mustroph *et al*., 2010; Reynoso *et al*., 2019; van Veen *et al*., 2016), our RNA-seq datasets indicate that the transcriptional reprogramming that accompanies the onset of waterlogging response in barley involves cellular processes such as carbohydrate metabolic processes, regulation of oxidative stress, and metal homeostasis. Many of the up and down-regulated genes identified in this study had also been shown to be differentially expressed in two other datasets generated using distinct barley varieties and different experimental conditions (Borrego-Benjumea *et al*., 2020; Luan *et al*., 2022; Luan *et al*., 2023). This overlap, together with the results of GO and RT-qPCR analyses validate our datasets. We also identified a set of 98 waterlogging response genes that were common to the five datasets generated in this study. Many of these common DEGs are homologs of ‘core hypoxia response genes’ that were identified across a range of plant species (Mustroph *et al*., 2009) and that are involved in processes such as cell wall metabolism, ethylene biosynthesis and signalling, carbohydrate metabolism and ROS regulation. Multiple transcription factors belonging to the ERF, bHLH, and WRKY families, which are known to be important for the onset of the waterlogging response program, are also among these 98 common DEGs. Hence, our data indicate that, as expected, the gene regulatory network that controls waterlogging response is conserved amongst monocots and dicots (Mustroph *et al*., 2010; Tamura and Bono, 2022).

Clustering analysis of the different DEGs identified groups of genes that behave similarly between the 5 varieties in response to waterlogging (*e.g*., all genes in clusters 14 and 9 are up or down-regulated, respectively). This analysis also revealed genes that are differentially expressed between the varieties in the absence of treatment (see clusters 1 and 16 (Figures 5F and 5H)). While their expression does not typically change in response to waterlogging, it is possible that either the low or high intrinsic expression of some of these genes may ‘predispose’ certain varieties to either sensitivity or tolerance to waterlogging. In other words, these genes could constitute variety-specific waterlogging susceptibility or tolerance factors. Validating this possibility would require (i) an in-depth characterization of the physiological responses and yield losses of a large number of varieties that would be first grouped based on the intrinsic expression level of genes of interest; (ii) dissecting the function of these genes, many of which have paralogs in barley; and (iii) generating barley plants that are mutated or that constitutively express these different genes, followed by a detailed characterization of the physiological response and yield under waterlogged conditions. Potential trade-offs on other traits of agronomic relevance and on resilience to other abiotic and biotic stresses would also need to be assessed. Considering the variety of gene functions present in clusters 1 and 16, it is however difficult to predict which genes (or gene families) are most likely to be of potential relevance to crop improvement. Their association with GWAS and QTL identification would most likely help to pinpoint the best candidates. For example, we compared our data to a list of 28 genes located in QTLs for waterlogging tolerance (Supp. Table S4) that were identified following a screen of nearly 700 barley varieties with a specific focus on root traits (i.e. the formation of adventitious roots and of root cortical aerenchyma) (Manik *et al*., 2022). A number of candidate genes that are a part of this list are differentially regulated between the control samples or the waterlogged samples of the different varieties we tested. These genes include the homolog of HRE2 (HORVU.MOREX.r3.6HG0621670) – one of the ERF-VII transcription factors that regulates the hypoxia response program, as well as a potassium transporter (HORVU.MOREX.r3.7HG0736590). This gene may be relevant because regulation of potassium flux during waterlogging has been shown to be important (Gill *et al*., 2018). Although the expression does not necessarily change in response to waterlogging, the intrinsic expression differences between varieties may be of interest to breeding waterlogging tolerant varieties in barley.

## Abbreviations

ACO: 1-aminocyclopropane-1-carboxylate oxidase 1-like
ADH: alcohol dehydrogenase
AGOUEB: association genetics of UK elite Barley
bHLH: basic helix-loop-helix
DEG: differentially expressed gene
ERF-VII: group VII ethylene response factor
GABA-T: γ-aminobutyric acid transaminase
GO: gene ontology
Golden P. /GP: Golden Promise
GWAS: genome-wide association studies
HB: hemoglobin
Hv: Hordeum vulgare
Inf: Infinity
MAPK: mitogen-activated protein kinase
NAC: NAM (no apical meristem), ATAF1 and −2, and CUC2 (cup-shaped cotyledon)
Pas: Passport
PCA: principal component analysis
PCO: plant cysteine oxidase
PDC: pyruvate decarboxylase
Pil: Pilastro
QTL: quantitative trait locus
RBOH: respiratory burst oxidase
Reg: Regina
ROS: reactive oxygen species
VST: variance stabilising transformations

## Acknowledgements

The authors are thankful to William Thomas for sharing seeds of the AGOUEB population, and to Susanne Barth and Carl Ng for helpful discussions. We also thank Daniel Daly for technical assistance with some of the waterlogging experiments

## Author contributions

AM, AJB and EG conducted experiments; AJB, AM, JB, FW and EG analyzed RNA-seq datasets and wrote the manuscript.

## Data availability

The datasets supporting the results of this article are included within the article (and its supplemental files). RNA-seq data have been deposited with the Gene Expression Omnibus (GEO) repository (at http://www.ncbi.nlm.nih.gov/) under GSE220532. *Note to reviewers*: the data has been submitted and the authors are waiting for publication before release.

