## Supplemental data for "Transcriptional analysis in multiple barley varieties identifies signatures of waterlogging response"

### Supplementary Tables

**Supplementary Table S1:** List of varieties used in this study. All varieties listed are winter varieties, except for *Golden Promise*, which is a spring variety. Apart from *Infinity* and *Golden Promise*, all varieties originated from the AGOUEB population (Thomas *et al.*, 2014).

| **Variety name** | **Row number** |
|---|---|
| *Golden Promise* | 2-row |
| *Infinity* | 2-row |
| *Passport* | 6-row |
| *Pilastro* | 6-row |
| *Regina* | 2-row |
| *Arma* | 6-row |
| *Isa* | 6-row |
| *Louise* | 2-row |
| *Retriever* | 2-row |
| *Dura* | 6-row |
| *Masquerade* | 2-row |
| *Madrigal* | 2-row |
| *Vesuvius* | 2-row |
| *Tapir* | 6-row |
| *Kosmos* | 6-row |
| *Maeva* | 6-row |
| *Siberia* | 6-row |
| *Breeze* | 2-row |
| *Cavalier* | 2-row |
| *Mahogany* | 2-row |
| *Tamaris* | 6-row |

**Supplementary Table S2:** List of oligonucleotides used in this study.

| Oligo name | Gene description | Gene ID | Oligonucleotide sequence (5' -> 3') | Reference |
|---|---|---|---|---|
| AM45 | HvACTIN | HORVU1Hr1G002840 | GCAAGTGGTCGTACTACTGGT ATCGTTC | (Miricescu *et al.*, 2021) |
| AM46 | HvACTIN | HORVU1Hr1G002840 | GGATCTTCATAAGGGAGTCCG TGAGAT | (Miricescu *et al.*, 2021) |
| AM8 | HvTUBULIN | HORVU.MOREX.r3.1 HG0082050 | TGGTCATTACACCATTGGCAA GGAGA | This study |
| AM9 | HvTUBULIN | HORVU.MOREX.r3.1 HG0082050 | GTGTATGTTGGGCGCTCAATG TCA | This study |
| AM49 | HvGAPDH | HORVU7Hr1G074690 | GCTCACTTGAAGGGTGGTGCC | This study |
| AM50 | HvGAPDH | HORVU7Hr1G074690 | TGATGGCATGAACAGTGGTCA TCAGAC | This study |
| HSP70F | HvHSP70 | HORVU.MOREX.r3.5 HG0526250 | GCTCAACATGGACCTCTTCAG G | (Walling *et al.*, 2018) |
| HSP70R | HvHSP70 | HORVU.MOREX.r3.5 HG0526250 | CCGACAAGGACAACATCATGG | (Walling *et al.*, 2018) |
| AM47 | HvADP | HORVU3Hr1G079700 | CCACCATCCCAACCATCGGTTT | This study |
| AM48 | HvADP | HORVU3Hr1G079700 | CCTGCGTATTCTGGAAGTAGT GCCT | This study |
| HvADH1F | HvADH1 | HORVU4Hr1G016810 | CACTGACCTGCCCAATGTC | (Mendiondo *et al.*, 2016) |
| HvADH1R | HvADH1 | HORVU4Hr1G016810 | GCACGCTGTGTGTGATGAA | (Mendiondo *et al.*, 2016) |
| AM51 | HvHB | HORVU4Hr1G066200 | CGGGAAGGAAGCCATGTCTGC | (Miricescu *et al.*, 2021) |
| AM52 | HvHB | HORVU4Hr1G066200 | TCTGCCTCGCCGACGG | (Miricescu *et al.*, 2021) |
| AM53 | HvPDC | HORVU4Hr1G056050 | CCGCCTACGAGAACTACAAGA GGATC | This study |
| AM54 | HvPDC | HORVU4Hr1G056050 | TTCATACCCGCAGCCCTCG | This study |

### Supplementary Figures

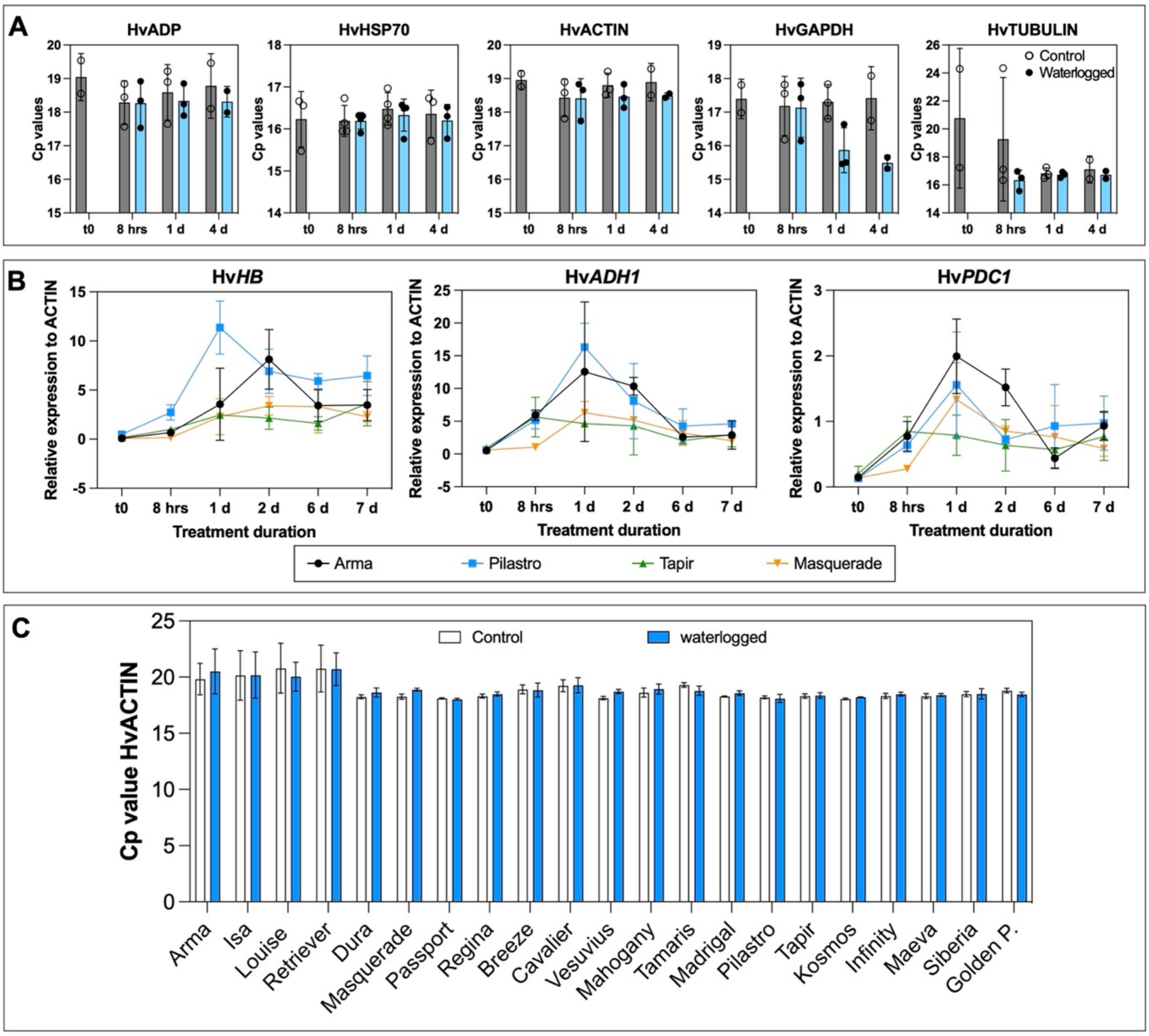


**Figure S1: Selection of reference genes for RT-qPCR analysis, and of time points for waterlogging experiments.** (A) Cp values for selected reference genes. Whole roots were collected at different time-points after the beginning of waterlogging treatment, and pooled for RNA extraction. Expression was determined using RT-qPCR. Cp values are shown for three biological replicates using the *Golden Promise* variety. (B) Transcriptional response of selected hypoxia response marker genes at different time points after the beginning of waterlogging treatment. Mean values of 2 biological replicates are shown. Error bars indicate standard deviations. (C) Mean Cp values for Hv*ACTIN* for all varieties and RT-qPCRs presented in Fig. 1. Error bars correspond to SEM.

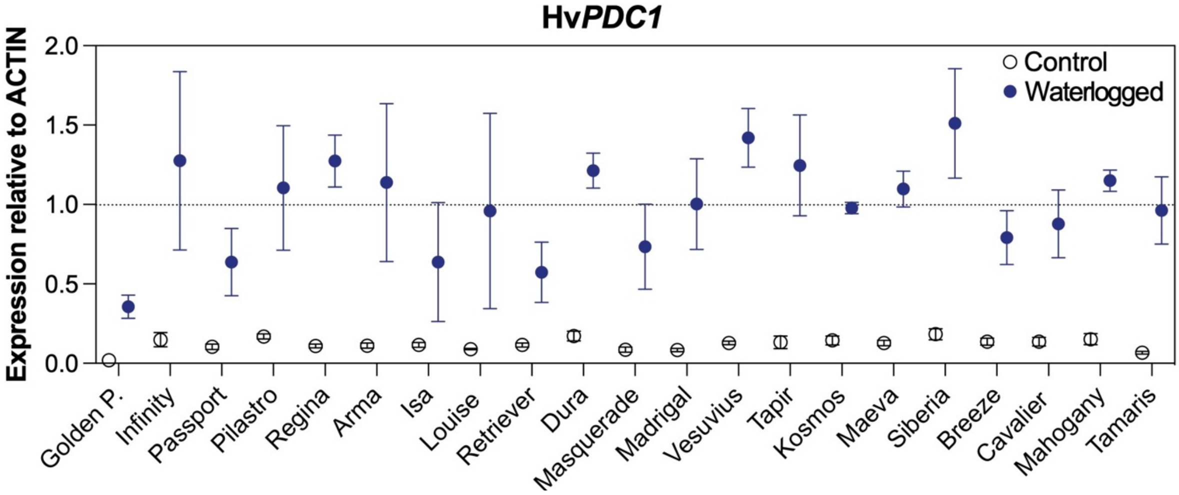


**Figure S2: Relative expression of Hv*PDC1* in selected varieties.** Hv*PDC1* expression relative to *ACTIN* in untreated (open symbols; normal watering) and waterlogged (blue symbols) plants after 24 h of treatment. The dashed line corresponds to the average relative expression of Hv*PDC1* for waterlogged samples for all varieties tested across all replicates. Data shown is from three biological replicates, except *Golden Promise* (*Golden P.*) which had 4 biological replicates. Each biological replicate corresponds to whole roots from 3 plants pooled. Error bars correspond to SEM. See Figure 1D for the results of two-way ANOVA.

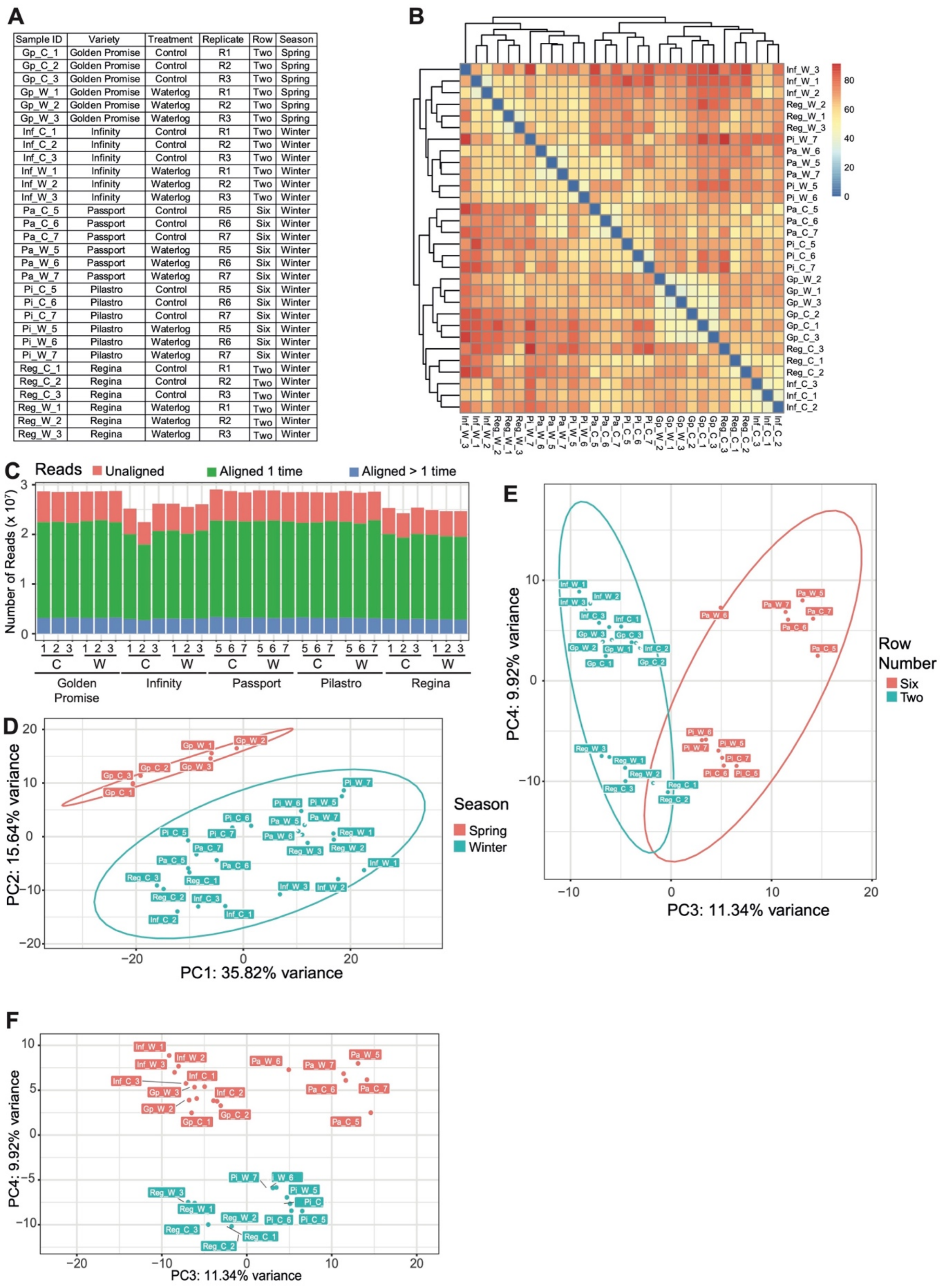

| Sample ID | Variety | Treatment | Replicate | Row | Season |
|---|---|---|---|---|---|
| Gp_C_1 | Golden Promise | Control | R1 | Two | Spring |
| Gp_C_2 | Golden Promise | Control | R2 | Two | Spring |
| Gp_C_3 | Golden Promise | Control | R3 | Two | Spring |
| Gp_W_1 | Golden Promise | Waterlog | R1 | Two | Spring |
| Gp_W_2 | Golden Promise | Waterlog | R2 | Two | Spring |
| Gp_W_3 | Golden Promise | Waterlog | R3 | Two | Spring |
| Inf_C_1 | Infinity | Control | R1 | Two | Winter |
| Inf_C_2 | Infinity | Control | R2 | Two | Winter |
| Inf_C_3 | Infinity | Control | R3 | Two | Winter |
| Inf_W_1 | Infinity | Waterlog | R1 | Two | Winter |
| Inf_W_2 | Infinity | Waterlog | R2 | Two | Winter |
| Inf_W_3 | Infinity | Waterlog | R3 | Two | Winter |
| Pa_C_5 | Passport | Control | R5 | Six | Winter |
| Pa_C_6 | Passport | Control | R6 | Six | Winter |
| Pa_C_7 | Passport | Control | R7 | Six | Winter |
| Pa_W_5 | Passport | Waterlog | R5 | Six | Winter |
| Pa_W_6 | Passport | Waterlog | R6 | Six | Winter |
| Pa_W_7 | Passport | Waterlog | R7 | Six | Winter |
| Pi_C_5 | Pilastro | Control | R5 | Six | Winter |
| Pi_C_6 | Pilastro | Control | R6 | Six | Winter |
| Pi_C_7 | Pilastro | Control | R7 | Six | Winter |
| Pi_W_5 | Pilastro | Waterlog | R5 | Six | Winter |
| Pi_W_6 | Pilastro | Waterlog | R6 | Six | Winter |
| Pi_W_7 | Pilastro | Waterlog | R7 | Six | Winter |
| Reg_C_1 | Regina | Control | R1 | Two | Winter |
| Reg_C_2 | Regina | Control | R2 | Two | Winter |
| Reg_C_3 | Regina | Control | R3 | Two | Winter |
| Reg_W_1 | Regina | Waterlog | R1 | Two | Winter |
| Reg_W_2 | Regina | Waterlog | R2 | Two | Winter |
| Reg_W_3 | Regina | Waterlog | R3 | Two | Winter |



**Figure S3: Assessing the quality of RNA-seq datasets.** (A) Details of samples used for RNA-Seq analysis. (B) Heatmap showing sample-to-sample Euclidean distance (generated by *PCA explorer* package in *R*). (C) Absolute number of reads aligned to Morex v3 for each sample. (D) PCA plot showing PC2 vs PC1 with ellipses grouping samples based on their sowing season. Plot is identical to Figure 3A, but highlights grouping based on PC2 instead of PC1.

(E) PCA plot showing PC4 vs PC3 with ellipses grouping by row number. (F) PCA plot showing PC4 vs PC3. Plot is identical to (E), but colour grouping based on PC4 instead of PC3.

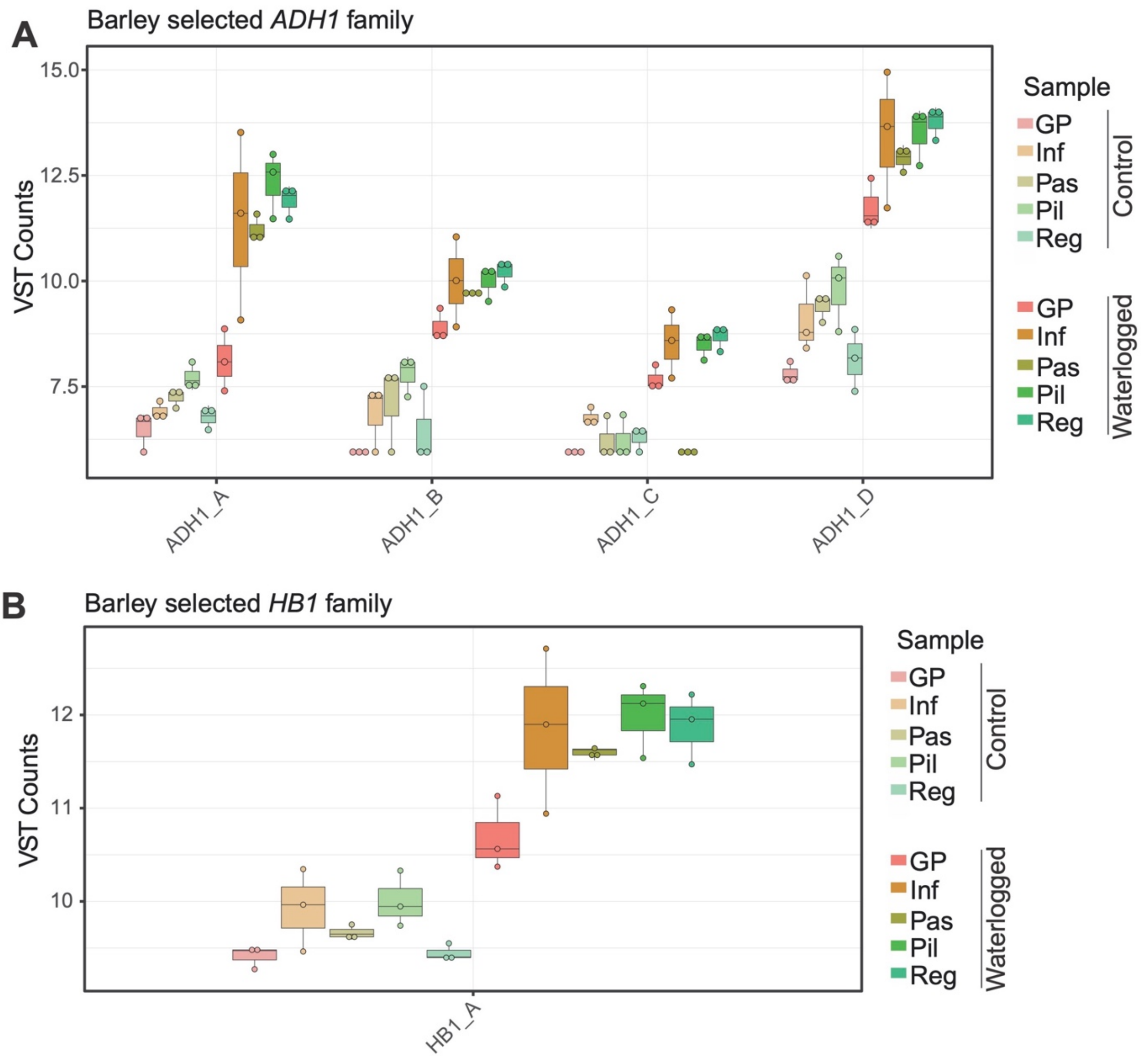


**Figure S4: Internal validation of RNA-seq datasets.** (A) Boxplots showing variance stabilised transformed (VST) read counts for barley orthologs of Arabidopsis *ADH1* that responded to waterlogging treatment: ADH1_A (HORVU.MOREX.r3.1HG0082960), ADH1_B (HORVU.MOREX.r3.2HG0115170), ADH1_C (HORVU.MOREX.r3.3HG0252910) and ADH1_D (HORVU.MOREX.r3.4HG0345740). (B) Boxplots showing VST read counts for a barley ortholog of Arabidopsis HB1 that responded to waterlogging treatment (HB1_A (HORVU.MOREX.r3.7HG0715010)). In (A) and (B), VST counts from control and waterlogged samples for *Golden Promise* (GP), *Infinity* (Inf), *Passport* (Pas), *Pilastro* (Pil) and *Regina* (Reg.) are shown.

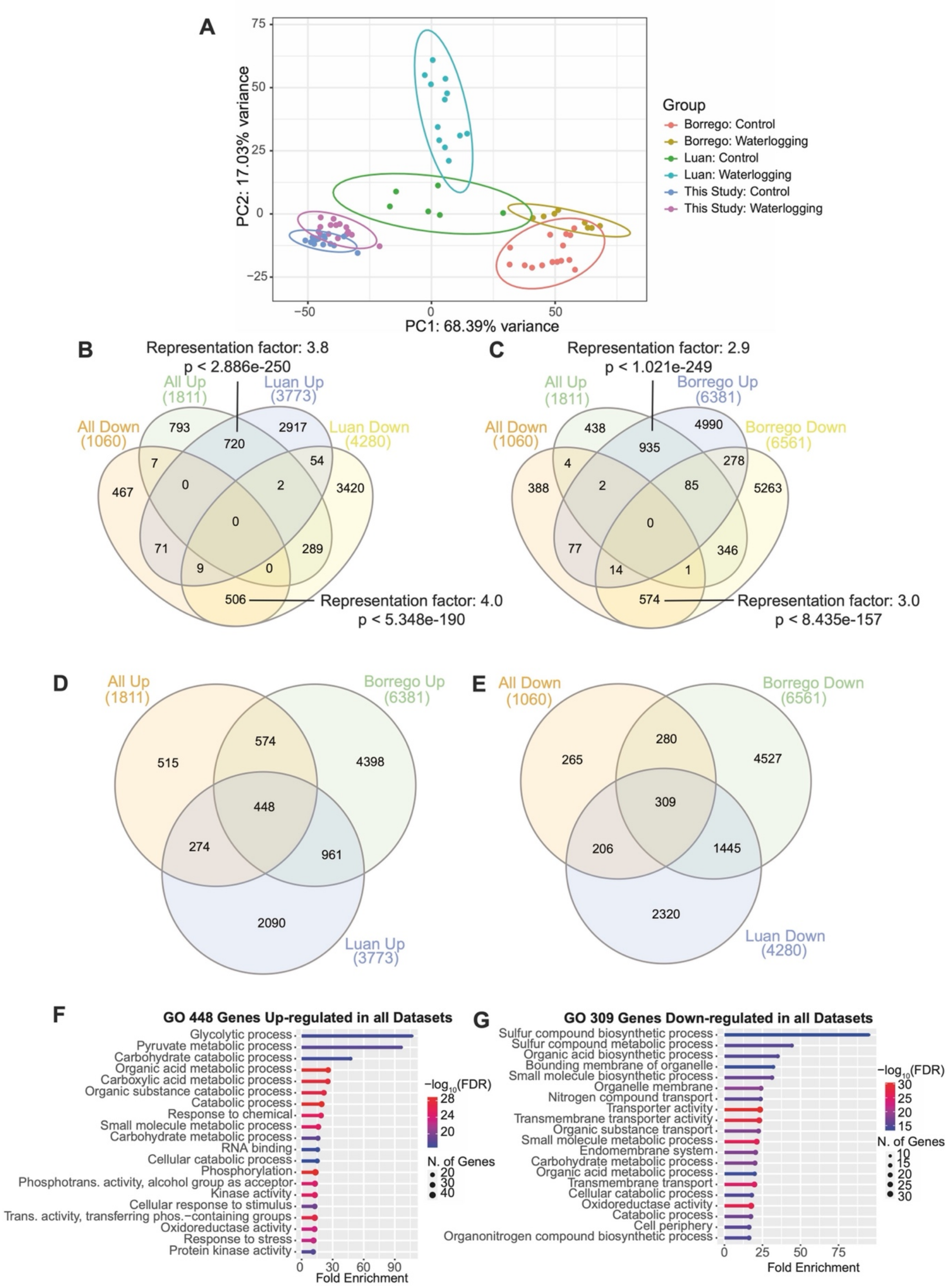


**Figure S5: Comparison of RNA-seq datasets with published datasets.** (A) PCA plot showing PC1 vs PC2 with ellipses grouping samples based on the dataset origin and treatment. (B) Overlap of DEGs identified in this study and those determined after re-analysis of raw data from Luan *et al.* (Luan *et al.*, 2022) (adjusted *p*-value < 0.05). (C) Overlap of DEGs identified in this study and those determined after re-analysis of raw data from Borrego-Benjumea *et al*. (Borrego-Benjumea *et al.*, 2020) (adjusted *p*-value < 0.05). (D) Overlap of up-regulated DEGs identified in this study with up-regulated DEGs from Luan et al. and Borrego-Benjumea et al. (E) Overlap of down-regulated DEGs identified in this study with down-regulated DEGs from Luan et al. and Borrego-Benjumea et al. (F) Top 20 GO terms enriched in 448 shared up-regulated DEGs identified in (D). (G) Top 20 GO terms enriched in 309 shared up-regulated DEGs identified in (E).

A

| Comparison | Group 1 | Group 2 |
|---|---|---|
| 1 | Golden Promise Control | Golden Promise Waterlog |
| 2 | Infinity Control | Infinity Waterlog |
| 3 | Passport Control | Passport Waterlog |
| 4 | Pilastro Control | Pilastro Waterlog |
| 5 | Regina Control | Regina Waterlog |
| 6 | Golden Promise Control | Infinity Control |
| 7 | Golden Promise Control | Passport Control |
| 8 | Golden Promise Control | Pilastro Control |
| 9 | Golden Promise Control | Regina Control |
| 10 | Infinity Control | Passport Control |
| 11 | Infinity Control | Pilastro Control |
| 12 | Infinity Control | Regina Control |
| 13 | Passport Control | Pilastro Control |
| 14 | Passport Control | Regina Control |
| 15 | Pilastro Control | Regina Control |
| 16 | Golden Promise Waterlog | Infinity Waterlog |
| 17 | Golden Promise Waterlog | Passport Waterlog |
| 18 | Golden Promise Waterlog | Pilastro Waterlog |
| 19 | Golden Promise Waterlog | Regina Waterlog |
| 20 | Infinity Waterlog | Passport Waterlog |
| 21 | Infinity Waterlog | Pilastro Waterlog |
| 22 | Infinity Waterlog | Regina Waterlog |
| 23 | Passport Waterlog | Pilastro Waterlog |
| 24 | Passport Waterlog | Regina Waterlog |
| 25 | Pilastro Waterlog | Regina Waterlog |

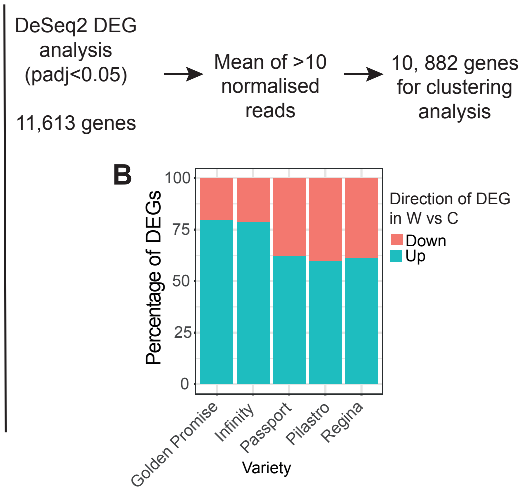


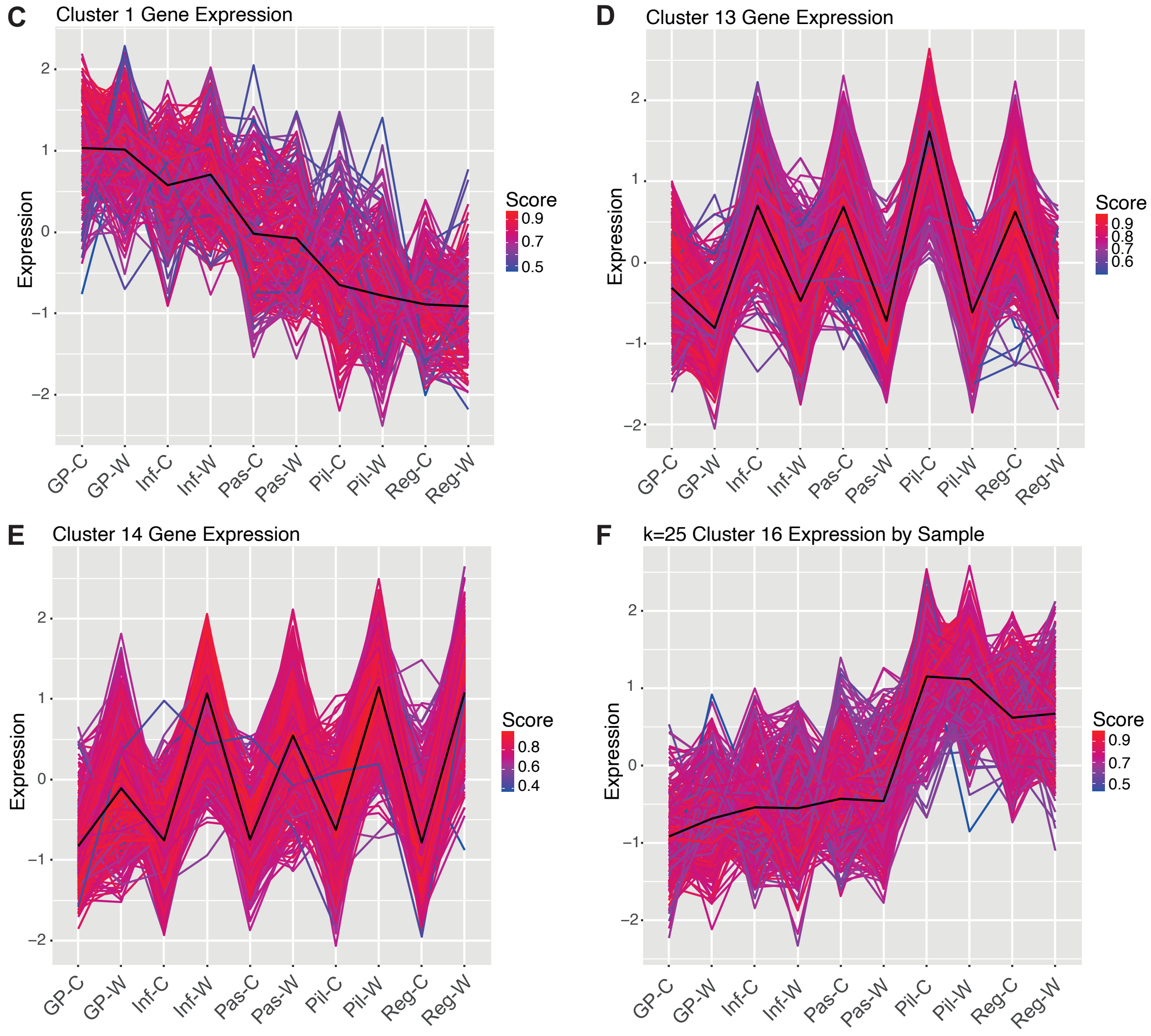


**Figure S6: Clustering Analysis.** (A) Structure of data used to generate DEG list used for *k*-means clustering analysis. The means of normalized read counts from 3 biological replicates generated from *DeSeq2* were filtered as follows. *Deseq2* DEG analysis was performed and results for 25 comparisons were extracted. Firstly, a list of all 11,613 differentially expressed genes (DEGs) filtered by adjusted *p*-value < 0.05 was generated by combining DEGs from waterlogged *vs* control samples from the same variety and DEGs from pairwise comparisons of each control to each other control variety and each waterlogged to each other waterlogged variety (25 DEG lists). Next the DEGs with a mean of <10 normalised reads were removed, leaving 10,882 genes for clustering analysis. (B) Number of up and down-regulated DEGs (filtered by adjusted *p*-value < 0.05) in waterlogged *vs* control samples for each variety as a percentage of all the DEGs in that sample. Gene expression patterns are shown for DEGs from cluster 1 (C), cluster 13 (D), cluster 14 (E) and cluster 16 (F). The expression of the core centroid is shown in black while the expression of each gene within the cluster is shown in colour. The score indicates the correlation of expression of each gene within a cluster to the expression pattern of the core centroid of that cluster. Scores closer to 1 indicate a better fit to the centroid expression pattern. Data is shown for control (C) and waterlogged (W) samples from *Golden Promise* (GP), *Infinity* (Inf), *Passport* (Pas), *Pilastro* (Pil) and *Regina* (Reg).
